# Leaf-to-whole plant spread bioassay for pepper and *Ralstonia solanacearum* interaction determines inheritance of resistance to bacterial wilt for further breeding

**DOI:** 10.1101/2021.01.27.428365

**Authors:** Ji-Su Kwon, Jae-Young Nam, Seon-In Yeom, Won-Hee Kang

## Abstract

Bacterial wilt (BW) disease by *Ralstonia solanacearum* is a serious disease and causes severe yield losses in chili peppers worldwide. Resistant cultivar breeding is the most effective in controlling BW. Thus, a simple and reliable evaluation method is required to assess disease severity and to investigate the inheritance of resistance for further breeding programs. Here, we developed a reliable leaf-to-whole plant spread bioassay for evaluating BW disease and then, using this, determined the inheritance of resistance to *R. solanacearum* in peppers. *Capsicum annuum* ‘MC4’ displayed a completely resistant response with fewer disease symptoms, a low level of bacterial cell growth, and significant up-regulations of defense genes in infected leaves compared to those in susceptible ‘Subicho’. We also observed the spreading of wilt symptoms from the leaves to the whole susceptible plant, which denotes the normal BW wilt symptoms, similar to the drenching method. Through this, we optimized the evaluation method of the resistance to BW. Additionally, we performed genetic analysis for resistance inheritance. The parents, F_1_ and 90 F_2_ progenies, were evaluated, and the two major complementary genes involved in the BW resistance trait were confirmed. These could provide an accurate evaluation to improve resistant pepper breeding efficiency against BW.

## 1. Introduction

Chili pepper (*Capsicum spp*.) is an important economic crop that belongs to the Solanaceae family alongside potatoes, tomatoes, and eggplants. Pepper is widely consumed as fresh, dried, or processed products and provides many essential vitamins, and capsaicin is used as a major spicy source in most global cuisines [1]. The consumption of pepper has increased in the last 40 years, with production ranging from 9 to approximately 41 million tons and the cultivation area increasing from 2.4 to approximately 3.8 million ha [2]. The world trade value of hot peppers has consistently increased during the last decade, with the second-largest quantity after the tomato in Solanaceae crops [3]. Pepper production is continuously challenged by biotic stresses such as fungi, viruses, and bacteria [4]. *Ralstonia solanacearum* is the causal agent of bacterial wilt (BW), one of the most destructive soil-borne bacterial pathogens in tropical and subtropical areas, with a wide host range of more than 400 plant species, especially the Solanaceae family including peppers [5]. BW by *R. solanacearum* is widely prevalent in peppers across much of Asia [6–8]. In China, that accounts for approximately half of the world’s production of peppers in 2017 (FAOSTAT), and the yield loss of BW from peppers is estimated to be approximately 20–50% in its cultivation area [9].

*R. solanacearum* species is divided into five races according to host range and five biovar according to the utilization of disaccharides and hexose alcohols [10]. *R. solanacearum* is also classified based on geographical origin: phylotype I from Asia, phylotype II from America, phylotype III from Africa, and phylotype IV from Indonesia [11]. Recently, a few studies have proposed to classify *R. solanacearum* into three species based on phylotype*: R. psedosolanacearum* (phylotype I and III), *R. solanacearum* (phylotype II), and *R. syzygii* (phylotype IV) [12, 13]. Thus, the *R. solanacearum* species complex includes phenotypically diverse and heterogeneous strains causing BW in a variable host range. This is one of the constraint factors of resistance studies on *R. solanacearum.* The pathogen can invade the plant through root wounds and subsequently resides in the xylem vessels to block water transport and ultimately kills the plant host [8, 14].

Most studies on resistance to *R. solanacearum* in plants have used two screening methods of *R. solanacearum,* such as root cut (soil)-drench and root-dipping inoculation [15-18]. However, both methods are difficult to determine the resistance degree according to the size of artificially root wounds lead to the standard deviation is large due to low uniformity after inoculation [17]. The stem-puncture inoculation method also has limitations as it is difficult to apply this approach depending on the crop [19]. The leaf-inoculation method by syringe is a commonly used method for bacteria inoculation, but this has not yet been reported to optimize a reliable bioassay in the resistance screening to *R. solanacearum* studies in peppers. This assay can infiltrate a relatively equal quantity of *R. solanacearum* into infected leaves and evaluate the quantification of pathogen growth in a plant. Additionally, leaf infiltration can recognize the inoculated leaves and non-inoculated systemic organs and establish disease scoring according to disease transmission in the whole plant.

To date, developed management programs of *R. solanacearum* are not sufficiently effective because chemical and biological controls are limited and ineffective in preventing the spread of *R. solanacearum* to the host plant [20, 21]. One of the most effective BW control methods is the development of a resistance cultivar in the crops. Presently, several resistance sources of BW resistance have been evaluated to develop resistant cultivars in *Capsicum spp.* Several pepper accessions have been reported among them, *C. annuum* ‘MC4’, *C. annuum* ‘MC5’, *C. annuum* ‘LS2341’, *C. annuum* ‘PBC473’, *C. annuum* ‘PBC 1347’, and *C. annuum* ‘PBC631’ are well known as the most strong BW resistant cultivars in various pathogens [22-24]. BW resistance is generally quantitatively inherited and is controlled by at least two genes in the pepper cultivar *C. annuum* ‘Mie-Midori’ [25]. Additionally, a pepper line *C. annuum* ‘PM687’ reported additive effects with two to five genes to control the BW resistance [26]. The pepper line *C. annuum* ‘LS2341’ is reportedly polygenic and linked to a major quantitative trait loci (QTL) named *Bw1* on chromosome 1 [27]. Recently, a major QTL named *qRRs-10.1* in chromosome 10 was revealed as a resistance pepper line *C. annuum* ‘BVRC1’ [28].

Among them, *C. annuum* ‘MC4’ is a well-known accession with a strong level of resistance to various *R. solanacearum* strains [15, 22, 29, 30]. However, despite reports of *C. annuum* ‘MC4’ resistance to BW, genetic inheritance analysis of BW resistance in *C. annuum* ‘MC4’ has not been determined yet because of pathogen strains complexity and a lack of an efficient bioassay of *R. solanacearum* in peppers. Here, we developed a fast and reliable bioassay for phenotype evaluation against *R. solanacearum* in pepper germplasms. Using this method, BW resistance and susceptible symptoms were distinctly confirmed, and we successfully detected disease symptoms through whole plant wilting and validation for pepper cultivars. Through this, a genetic inheritance analysis of BW resistance was investigated in the parents, F_1_ and F_2_ progeny populations. The BW resistance trait in ‘MC4’ confirmed to be affected with at least two major complementary genes.

## 2. Results

### 2.1. Identification of leaf wilt symptoms between resistant and susceptible pepper

To identify the response of pepper plants on leaf wilting by *R. solanacearum*, we performed an infiltration of *R. solanacearum* SL1931 (hereafter SL1931) with 10^6^ CFU/mL in resistant ‘MC4’ and susceptible ‘Subicho’ to BW. We observed phenotypes of the infiltrated area for both cultivars from day 1 to day 4 after inoculation. Disease symptoms, leaf wilting, and yellowing with necrosis were observed in ‘Subicho’ at 3 days after inoculation (dai), whereas ‘MC4’ displayed no symptoms within 4 dai (Fig. 1A). To confirm the resistant response between ‘MC4’ and ‘Subicho’, we quantified the level of bacterial cell growth in both cultivars. The differences in bacterial growth were observed at 2 dai but were significant from 3 to 5 dai, displaying 10 to 100 times more bacterial growth in ‘Subicho’ than in ‘MC4’ (Fig. 1B).

**Figure 1.**
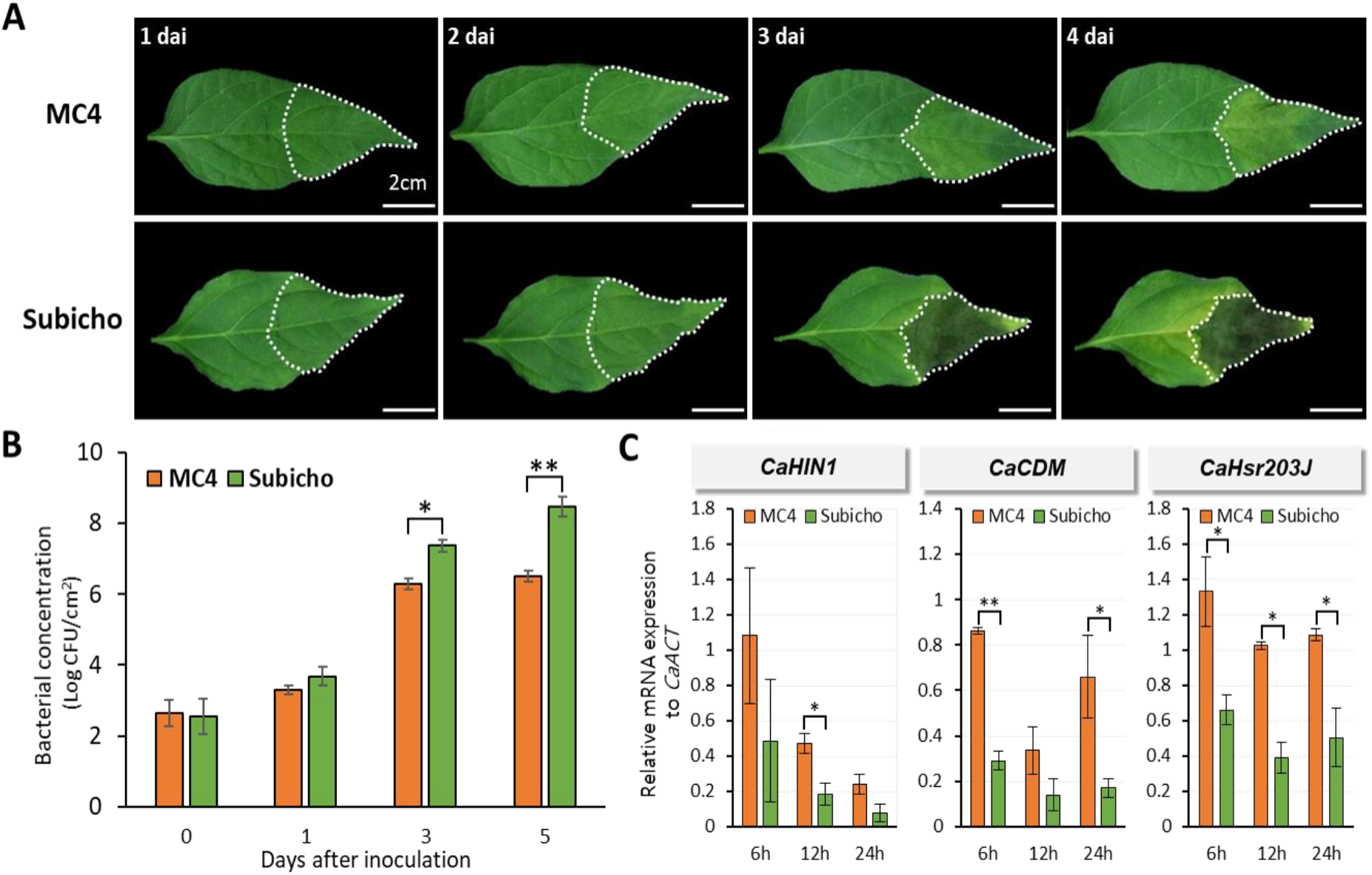
Assessment of BW response by *R. solanacearum* in pepper leaves. The eight-leaf stage seedlings were inoculated with *R. solanacearum* SL1931 by leaf infiltration with bacterial suspensions 1 × 10^6^ CFU/mL to give inoculum volume of 0.1mL per leaf. The plants were incubated in a growth room at 28°C with 16-hour light a day. (A), Difference of necrotic lesions present in the leaf of inoculated ‘MC4’ and ‘Subicho’. The symptom of ‘MC4’ (R) and ‘Subicho’ (S) leaf according to 1, 2, 3 and 4 days after inoculation (dai) is shown. (B) Bacterial multiplication in the apoplast of ‘MC4’ and ‘Subicho’ leaves. Bacterial suspension is 1 × 10^4^ CFU/mL to give inoculum volume of 0.1 mL/leaf. Total six to eight leaves used one experiment. Each vertical bar represents the S.E from two independent experiment. (C) Reverse-transcription polymerase chain reaction of defense-related expression gene colonization levels in ‘MC4’ and ‘Subicho’ against *R. solanacearum.* A graph represent the difference of relative expression in ‘MC4’ (R), ‘Subicho’ (S) leaves according to cell death marker in 6h and 12h after inoculation. Asterisks indicate statistically significant differences in 5 dai according to Student’s t-test (\**p < 0.05*, \*\**p<0.01*).

Although no differences were observed during infection until 3 dai, the resistant response of *R. solanacearum*-inoculated leaves changed dramatically within a day between the two pepper cultivars (Fig. 1C). We measured the transcript expression of cell-death related genes, *CaHIN1*, *CaCDM*, and *CaHsr203J,* that were expressed during the resistant response with hypersensitive response (HR)-like cell death induced by various pathogens [31-33]. The expression level of the *CaHIN1* gene was significantly increased in ‘MC4’ than in ‘Subicho’ at 12 h after inoculation (hai), and the *CaCDM* gene was also significant at 6 and 24 hai. Additionally, we confirmed the transcript expression levels of the *CaHsr203J* gene was significantly increased in ‘MC4’ than in ‘Subicho’ at all three-time points (Fig. 1C). Collectively, these data indicated that ‘MC4’ also has a suitable resistance to leaf wilting disease by *R. solaneacerum* alongside BW disease through root infection [15].

### 2.2. BW symptoms by *R. solanacearum* through leaf-to-whole plant spread bioassay (LWB)

To further understand the spectrum of defense responses to BW disease, the difference in phenotype of whole plants after leaf infection in the two cultivars was observed during 15 dai (Fig 2). ‘Subicho’ started to display wilt disease symptoms with the injected leaf abscising at 5 dai, whereas no differences in ‘MC4’ were observed until 10 dai. On the 15 dai, ‘MC4’ had a symptom of shedding and/or yellowing only inoculated leaves while ‘Subicho’ had wilted and the whole plant died, which is a common BW disease symptom (Fig. 2A and 2B). We confirmed the same wilt symptoms as the soil (root)-drenching inoculation method, although the leaf infection was conducted. We also represent the wilting rate (%) data that analyzed two replicate experiments using 30 plants for each cultivar (Fig. 2C). With consistency, ‘Subicho’ started to wither 6 dai, and rapid wilting progressed until 10 dai, and almost all the plants died on the 15 dai. Conversely, the ‘MC4’ was healthy with no wilting symptoms until two weeks after inoculation. Collectively, through the LWB, we could demonstrate quantified resistance and susceptible phenotypes to BW disease (Fig. 2C).

**Figure 2.**
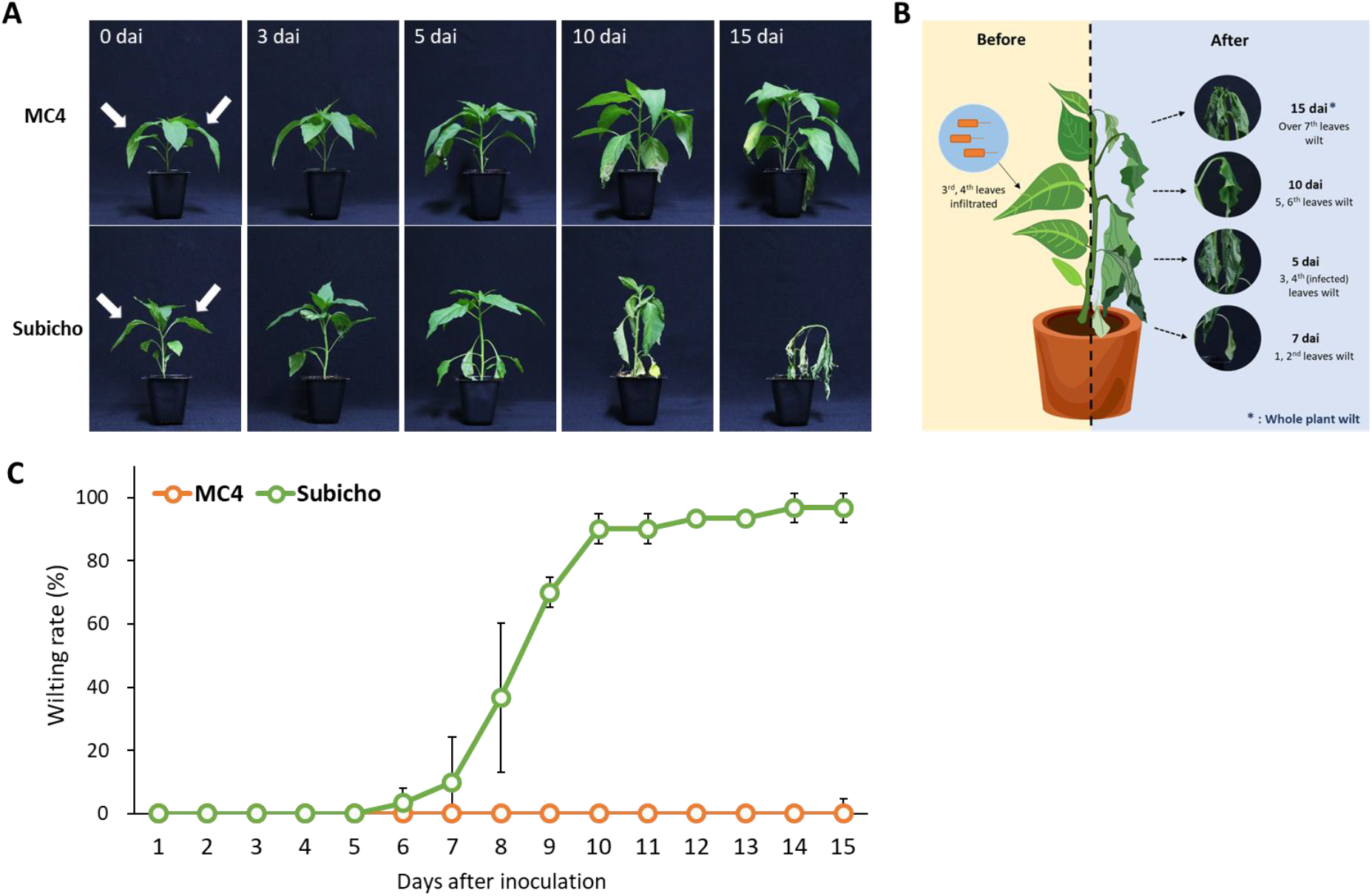
The difference in disease symptoms of leaf to whole plant spread bioassay (LWB) in pepper. Three week after transplanting, the eight-leaf stage seedlings were inoculated with *R. solanacearum* SL1931 by leaf infiltration with bacterial suspensions 1 × 10^6^ CFU/mL to give inoculum volume of 0.1 mL/leaf. (A), Difference of bacterial wilt (BW) symptom progression in inoculated ‘MC4’ and ‘Subicho’. The phenotype of ‘MC4’ (R), ‘Subicho’ (S) according to 0, 3, 5, 10 and 15 dai is shown. (B), Illustration of a procedure in which the whole plant withers after leaf-infiltration. (C), Progress degree of wilt disease on ‘MC4’ and ‘Subicho’. Disease severity of the plants was investigated every day after leaf inoculation. Green and Orange lines indicate ‘Subicho’ and ‘MC4’. In total, 30 plants were analyzed for each cultivar. The arrows show inoculated leaves. Each data point represents the mean disease index for two independent experiment.

### 2.3 Development of an efficient evaluation system for resistance to *R. solanacearum* in pepper

A clear score criterion for resistant evaluation was established on the DSI from 0 to 4 using LWB, which demonstrated identical BW symptoms with other methods (Fig. 3A-E) [28, 34]. Additionally, we measured to closely examine the abscission of leaves in the stem after wilting (Fig. 3F-H). A score of one of the DSI represents 3rd, 4th leaf abscission that is injected leaves simultaneously, the wilt of 2 leaves stands for 25% wilt symptoms (1 score of DSI) in total 8-leaf stage (Fig. 3B, F). The DSI of 2 scores designated when three or/and four leaves wither or abscission which is a symptom of 50% wilt in 8-leaf-stage (Fig. 3C, G). The degree of more than half of the leaves wilted and a few alive is determined as DSI of 3 scores (Fig. 3D, H). A plant with a DSI of < 2 was considered resistant (R), 2 ≤, a DSI of < 3 was moderate resistance (MR), and susceptible (S) was defined as a DSI of ≥ 3 in 15 dai based on Fig. 2A and 2C results.

**Figure 3.**
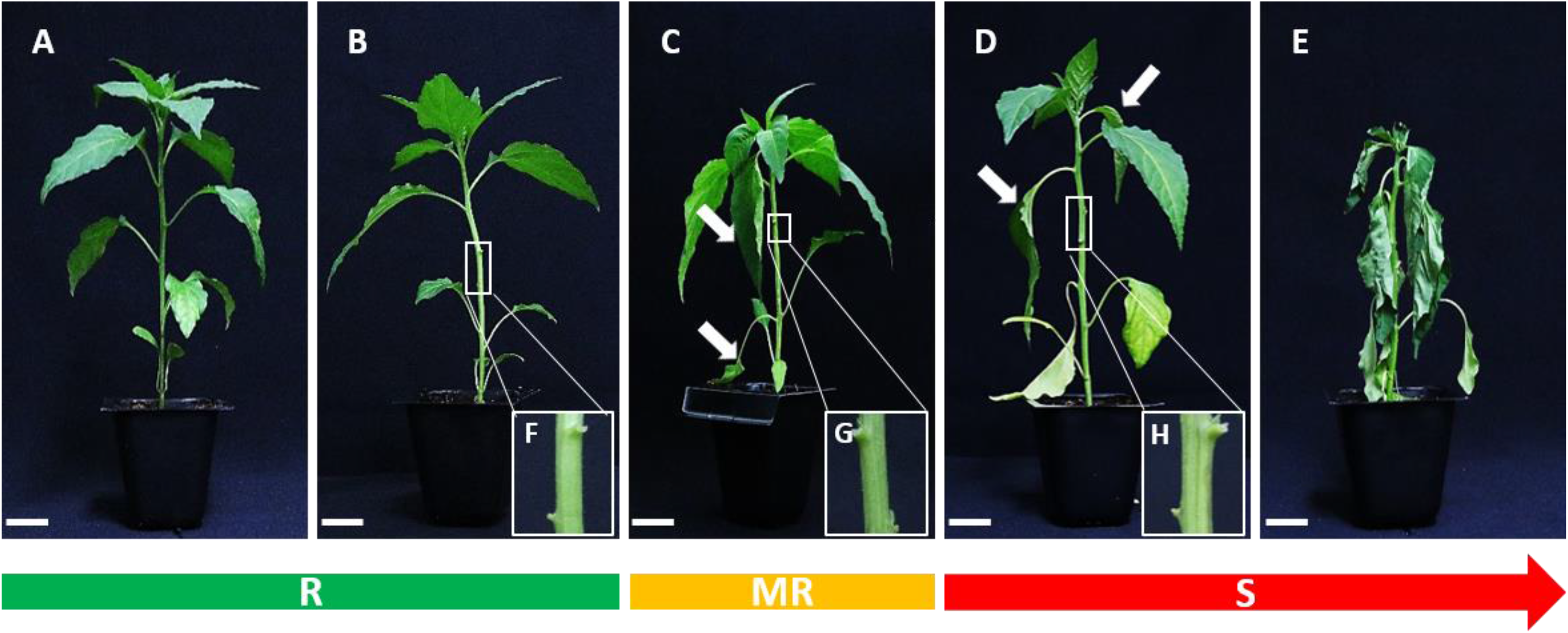
The disease symptoms scale ranging from 0 to 4 for BW evaluation. The (A) to (E), photo represent 0 (no symptoms) to 4 (complete wilting) wilt symptoms stages. The BW phenotype of three stages denoted resistance with a green bar, moderate resistance with a yellow bar, and susceptible with a red bar. The white arrows indicate wilt and abscission leaves. The white under bar signifies 2cm.

Next, to ensure the optimal evaluation for BW resistance in peppers, we determined the optimal conditions of LWB. Among the environmental conditions, temperature most affects the vitality of *R. solanacearum* that inhabits tropical and subtropical areas. Appropriate temperature conditions (28 – 32 °C) of screening for bacterial wilt have been identified in several studies on various crops and *R. solanacearum* strains [17, 35, 36]. We followed the above temperature and plant growth conditions and experimented to confirm the suitable inoculum concentration. Here, we compared four inoculum concentration levels from 10^3^ CFU/mL to 10^6^ CFU/mL at 10-fold intervals (Fig. 4). Differences in BW symptoms between the two cultivars can be verified at all concentrations of 10^3^ CFU/mL to 10^6^ CFU/mL according to statistical analysis. The DSI of 10^3^ CFU/mL concentration scored an average 2.6 in 20 dai, which does not represent a completely susceptible phenotype, and we considered it unsuitable. In the case of 10^4^ CFU/mL, the disease progression was similar with 10^3^ CFU/mL until 11 dai, , and after that disease progression was similar with 10^5^ CFU/mL from the 15 to 20 dai. The 10^6^ CFU/mL concentration was represented as the most suitable result. Resistance in ‘MC4’ maintained a DSI score of less than 1, whereas ‘Subicho’ displayed a fast-wilting symptom that scored a mean value of 3.8 until 20 dai (Fig. 4). The 10^6^ CFU/mL concentration displayed relatively quick and clear phenotypic differences between resistant and susceptible cultivars than others at 10 dai, and the condition was maintained until 20 dai.

**Figure 4.**
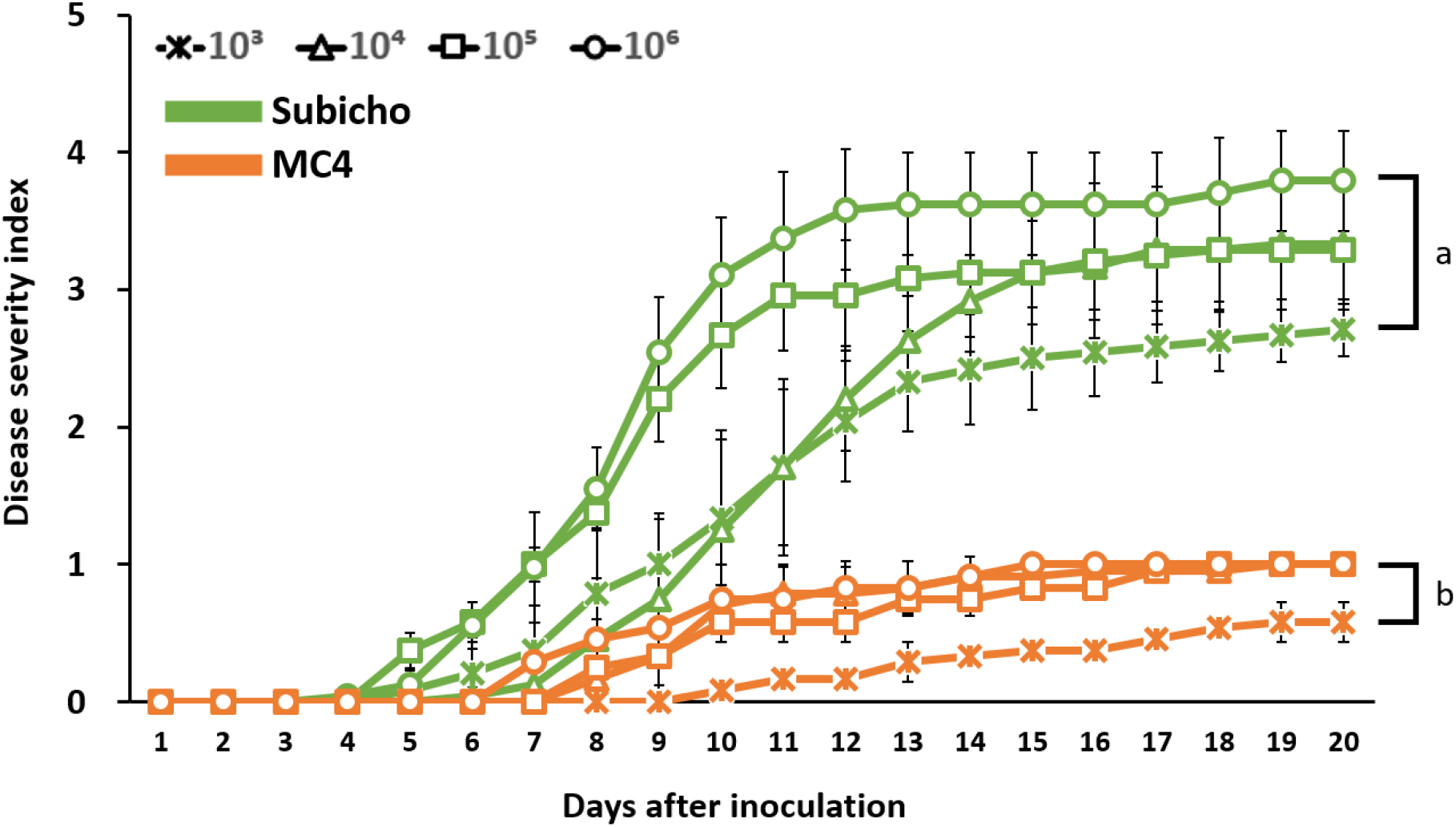
Occurrence of bacterial wilt on seedling of two pepper cultivars according to inoculum concentration. Three week after transplanting, the eight-leaf stage seedlings were inoculated with *R. solanacearum* SL1931 with bacterial suspensions (1 × 10^3^, 1 × 10^4^, 1 x 10^5^ and 1 × 10^6^ CFU/mL) to give inoculum volume of 0.1 mL/leaf. Disease severity of the plants was investigated every day after inoculation. Green and Orange lines indicate ‘Subicho’ and ‘MC4’. Each bar represents the S.E from three independent experiment with 24 plants. Values in the labeled with the same letter within each inoculum concentration are not significantly different in Duncan’s multiple range test at *P = 0.05.*

To further confirm and validate the LWB method, 12 commercial cultivars were re-evaluated for resistance to *R. solanacearum.* The DSI of BW symptoms was checked daily according to LWB (Fig. 5 and Table 1), which displayed R, MR, and S groups. We observed that ‘PR-Daedeulbo’ and ‘Supermanidda’ wilt in most individuals scored 3.3 and 3.9, respectively, of which ‘Supermanidda’ is as susceptible as ‘Subicho’ (Fig. 5). ‘Suppermanidda’ started to wilt early at 4 dai, also its disease progression is similar to ‘Subicho’, an S-control cultivar. ‘PR-Daedeulbo’ was a MR phenotype until 14 dai, but then exceeded a score of 3 with over 70% of individuals dead and was thus identified as an S cultivar. By contrast, ‘PR-Jangwongeunje’ and ‘PR-Chengyang’ belonged to the resistance category with the same DSI score of 1.8 in 20 dai but did not display the resistance of ‘MC4’ (0.6 score). The other 8 pepper accessions were denoted MR with scores between 2.0 to 2.5, and a wilt rate (%) at approximately half of the total tested plants for each (data not presented).

**Figure 5.**
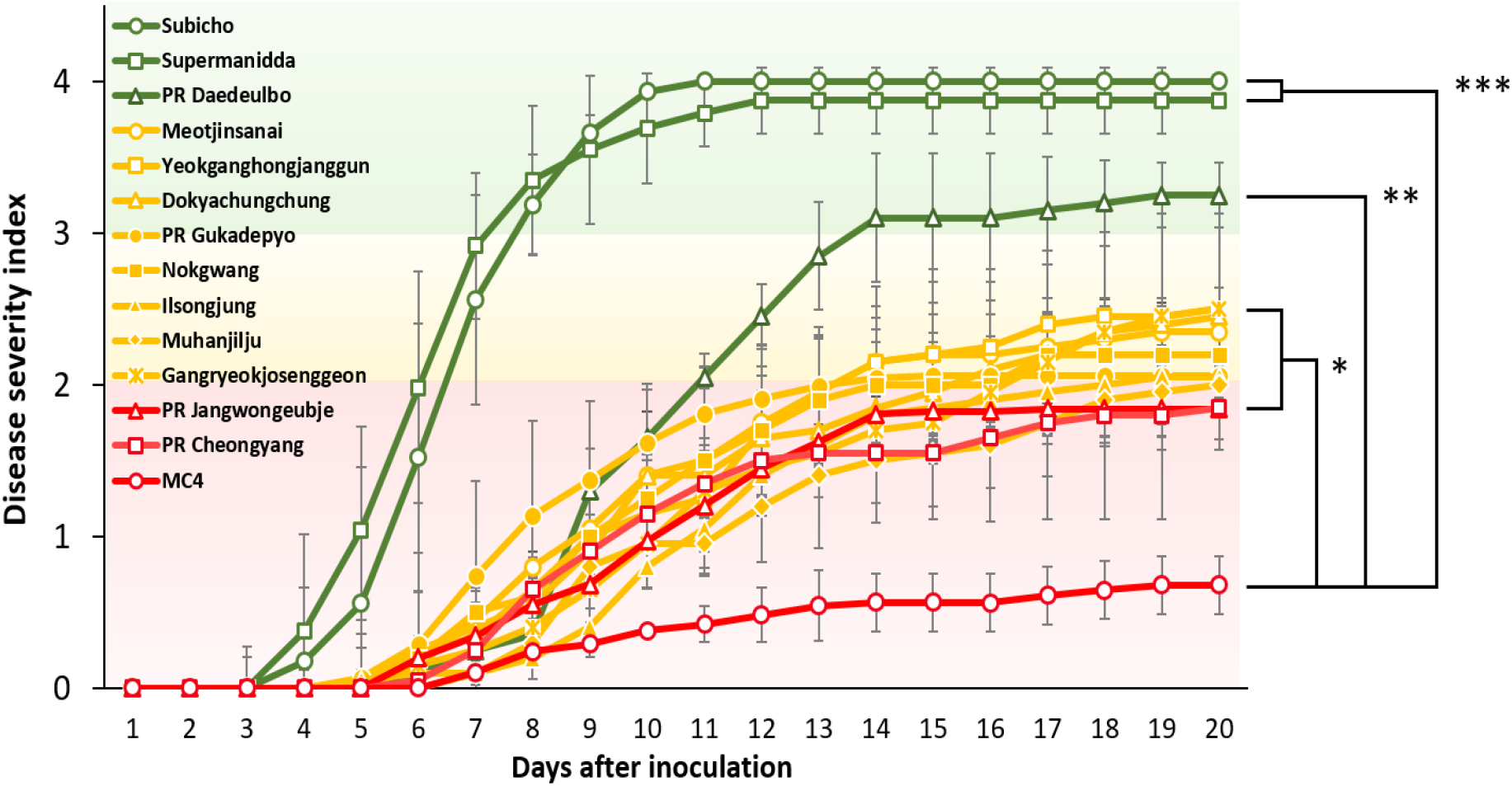
Disease progression through leaf to whole plant spread bioassay (LWB) in 12 pepper accessions. The eight-leaf stage seedlings were inoculated with *R. solanacearum* SL1931 with bacterial suspensions 1 × 10^6^ CFU/mL to give inoculum volume of 0.1 mL/leaf. A line graph area of red, yellow and green indicated resistance (R), moderate resistance (MR), susceptible (S) and, the color of line was expressed the same as the areas based on the DSI score of the bacterial wilt on 20 dai for each cultivar. Each data point represents the mean disease index from at least two independent experiments. Each bar represents the S.E from three independent experiment with 24 plants. Asterisks indicate statistically significant differences (\**p < 0.05*, \*\**p < 0.01*, \*\*\**p < 0.001*) in AUDPC (0 to 15d) according to Student’s t-test with ‘MC4’.

**Table 1.**
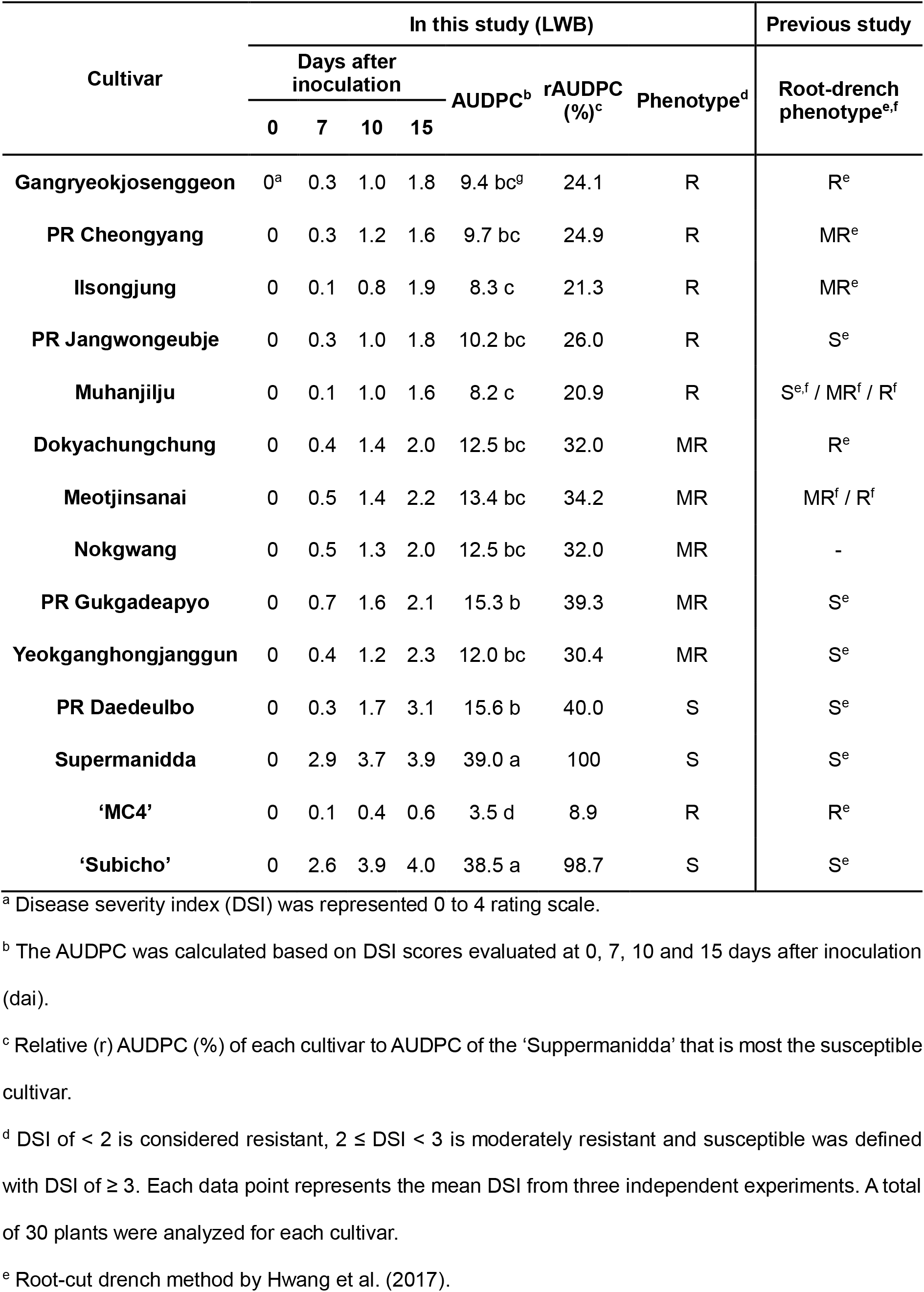
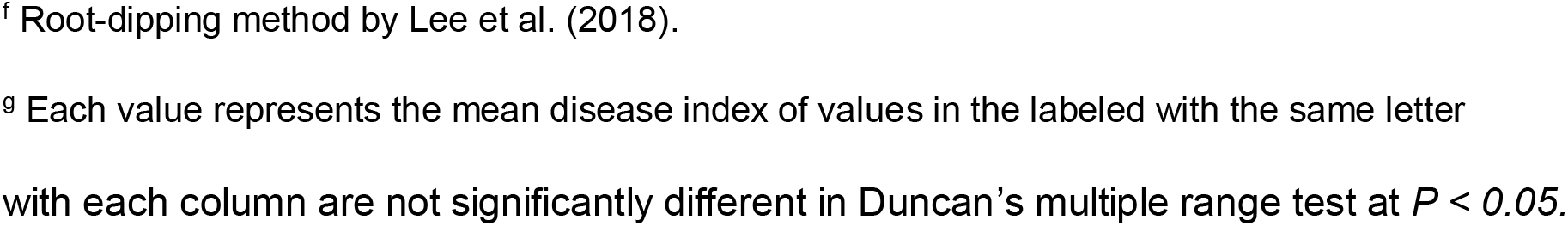
BW phenotype of LWB in 12 commercial chili pepper cultivars to *R. solanacearum*

| Cultivar | In this study (LWB) |  |  |  |  |  |  | Previous study |
| --- | --- | --- | --- | --- | --- | --- | --- | --- |
|  | Days after inoculation |  |  |  | AUDPC<sup>b</sup> | rAUDPC (%)<sup>c</sup> | Phenotype<sup>d</sup> | Root-drench phenotype<sup>e,f</sup> |
|  | 0 | 7 | 10 | 15 |  |  |  |  |
| Gangryeokjosenggeon | 0<sup>a</sup> | 0.3 | 1.0 | 1.8 | 9.4 bc<sup>g</sup> | 24.1 | R | R<sup>e</sup> |
| PR Cheongyang | 0 | 0.3 | 1.2 | 1.6 | 9.7 bc | 24.9 | R | MR<sup>e</sup> |
| Ilsongjung | 0 | 0.1 | 0.8 | 1.9 | 8.3 c | 21.3 | R | MR<sup>e</sup> |
| PR Jangwongeubje | 0 | 0.3 | 1.0 | 1.8 | 10.2 bc | 26.0 | R | S<sup>e</sup> |
| Muhanjilju | 0 | 0.1 | 1.0 | 1.6 | 8.2 c | 20.9 | R | S<sup>e,f</sup> / MR<sup>f</sup> / R<sup>f</sup> |
| Dokyachungchung | 0 | 0.4 | 1.4 | 2.0 | 12.5 bc | 32.0 | MR | R<sup>e</sup> |
| Meotjinsanai | 0 | 0.5 | 1.4 | 2.2 | 13.4 bc | 34.2 | MR | MR<sup>f</sup> / R<sup>f</sup> |
| Nokgwang | 0 | 0.5 | 1.3 | 2.0 | 12.5 bc | 32.0 | MR | - |
| PR Gukgadeapyo | 0 | 0.7 | 1.6 | 2.1 | 15.3 b | 39.3 | MR | S<sup>e</sup> |
| Yeokganghongjanggun | 0 | 0.4 | 1.2 | 2.3 | 12.0 bc | 30.4 | MR | S<sup>e</sup> |
| PR Daedeulbo | 0 | 0.3 | 1.7 | 3.1 | 15.6 b | 40.0 | S | S<sup>e</sup> |
| Supermanidda | 0 | 2.9 | 3.7 | 3.9 | 39.0 a | 100 | S | S<sup>e</sup> |
| ‘MC4’ | 0 | 0.1 | 0.4 | 0.6 | 3.5 d | 8.9 | R | R<sup>e</sup> |
| ‘Subicho’ | 0 | 2.6 | 3.9 | 4.0 | 38.5 a | 98.7 | S | S<sup>e</sup> |

<sup>a</sup> Disease severity index (DSI) was represented 0 to 4 rating scale.
<sup>b</sup> The AUDPC was calculated based on DSI scores evaluated at 0, 7, 10 and 15 days after inoculation (dai).
<sup>c</sup> Relative (r) AUDPC (%) of each cultivar to AUDPC of the ‘Supermanidda’ that is most the susceptible cultivar.
<sup>d</sup> DSI of < 2 is considered resistant, 2 ≤ DSI < 3 is moderately resistant and susceptible was defined with DSI of ≥ 3. Each data point represents the mean DSI from three independent experiments. A total of 30 plants were analyzed for each cultivar.
<sup>e</sup> Root-cut drench method by Hwang et al. (2017).
<sup>f</sup> Root-dipping method by Lee et al. (2018).
<sup>g</sup> Each value represents the mean disease index of values in the labeled with the same letter
with each column are not significantly different in Duncan's multiple range test at $P < 0.05$ .

Furthermore, on the LWB method, we compared the BW phenotype with the previous root and soil inoculation methods (Table 1). We also calculated the area under the disease progress curve (AUDPC) and relative (r) AUDPC (%) based on DSI scores at 7, 10, and 15 dai. Not only the DSI for wilting evaluation, but also the rAUDPC (%) value was able to distinguish between 0–30 % R, 30–40% MR, and 40–100% as S 15 dai [15]. The AUDPC and rAUDPC (%) were distributed as 3.5 and 8.9% in ‘MC4’, and ‘Subicho’ was 38.5 and 100%, displaying significant results as controls. Of the 12 commercial pepper cultivars, the rAUDPC (%) of ‘Supermanidda’ (100%) and ‘Muhanjilju’ (20.9%) had greater results or BW susceptibility and resistance, respectively. We compared the traits with the other inoculation methods and analyzed the DSI score of the BW phenotype 15 dai when ‘Subicho’ was in a saturating state. The ‘Gangryeokjosenggeon’ (R), ‘Meotjinsanai’ (MR), ‘PR-Daedeulbo’ (S), and ‘Supermanidda’ (S) have the same traits in either inoculation method (Table 1). However, the traits of the root-drenching method in ‘PR-Cheongyang’, ‘Ilsongjung’, ‘Muhanjilju’, and PR-‘Jangwongeubje’ were MR or S phenotypes [15], but in this study represented all R phenotypes. ‘Muhanjilju’ and ‘Meotjinsanai’ also displayed previously different traits with S, MR, and R on infection methods and/or *R. solanacearum* strains [16], whereas we observed R and MR uniformly in each cultivar, respectively (Table 1 and Fig. 5). Even though it could be difficult to determine the exact traits to BW, our results suggested that the LWB could be a simple and reliable evaluation method for BW resistant screening in peppers.

### 2.4 Inheritance analysis of resistance to *R. solanacearum* in pepper

To analyze the inheritance of resistance to *R. solanacearum* in ‘MC4’, the parents, F_1_ and F_2_, progenies were evaluated until the disease progressed at 30 dai (Table 2 and Table S2, Fig. 6). The parents, ‘MC4’ and ‘Subicho’, maintained resistance and susceptibility, respectively. The wilting progression of F_1_ plants was conspicuously slower than in the susceptible parent, and the wilt rates of F_1_ until 20 dai were closer to the resistant parent. In generation F_2_, the individuals were distributed on most DSI scores, but resistant plants were most common both at 15 and 20 dai. However, these BW symptoms in parents, F_1_ and F_2_, developed continuously until the end of the experiment at 30 dai (Fig. 6 and Table 2). These results suggested that BW resistance acts as a QTL with a few genes in ‘MC4’.

**Table 2.**
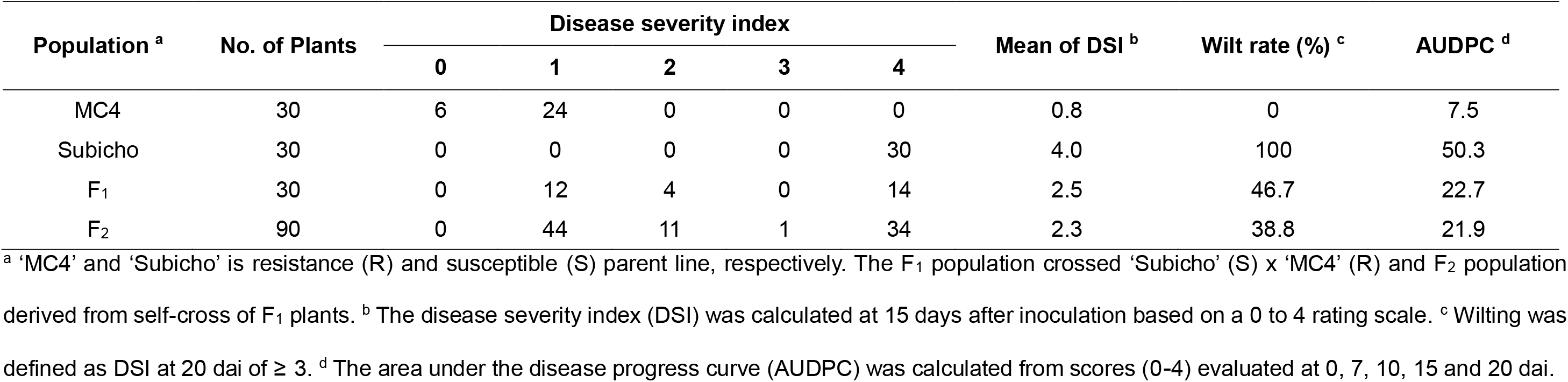
Disease evaluation design and the number of plants to parents and their progenies based on disease severity index in 20 dai against *R. solanacearum* SL1931 strain.

**Figure 6.**
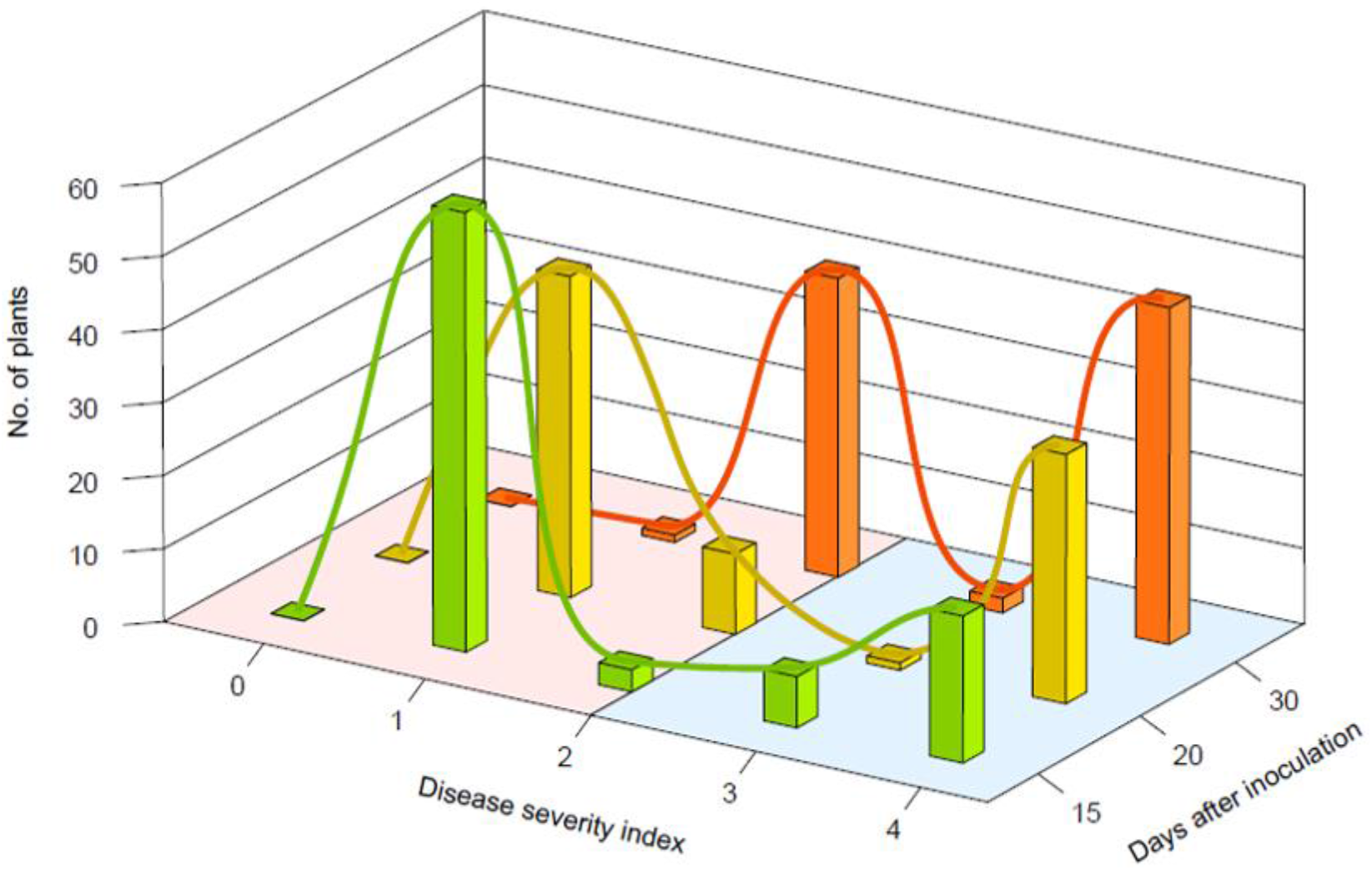
Histograms and curve graphs represented the number of plants’ phenotype segregation based on disease severity scores of the F_2_ population (n=90) at 15 (green bar), 20 (yellow bar), and 30 dai (orange bar). The plants were inoculated with *R. solanacearum* strain SL1931, the bacterial suspensions are 1 x 10^6^ CFU/ml to give inoculum volume of 0.1 mL/leaf at fully expended 3rd and 4th leaf stages in a plant. The red zone and blue zone represented resistance and susceptible, respectively.

We measured the segregation ratio of BW resistance with the chi-square analysis in the F_2_ population with disease progression. At 15 dai, segregation in F_2_ yielded 63 resistant and 27 susceptible plants that fitted closely to a 11:5 ratio (P > 0.5) and 3:1 ratio (P > 0.1). It appeared more closely at an 11:5 ratio than 3:1, which demonstrated that BW resistance was predominantly controlled by at least one major factor and/or two major alleles around two weeks after inoculation. At 20 dai, resistant plants in the F_2_ prevailed with 61 resistant plants versus 29 susceptible, which nearly matched a 9:7 ratio (P > 0.5) and 11:5 ratio (P > 0.1). Lastly, the segregation was represented as a 9:7 ratio (P > 0.05) with 42 resistant plants versus 48 susceptible at 30 dai (Fig. 6 and Table 3). According to these chi-square tests, there were significant differences in the segregation ration during pepper-*R. solanacearum* interaction. The BW resistance in ‘MC4’ may be affected by a major dominant factor until 15 dai alongside at least two factors controlling the resistance after the 20 dai. Additionally, the separation ratios of 11:5 and 9:7 were consistently represented with a high p-value closest at 20 and 30 dai, which indicated that two complementary dominant genes could mainly control the resistance to BW in ‘MC4’.

**Table 3.**
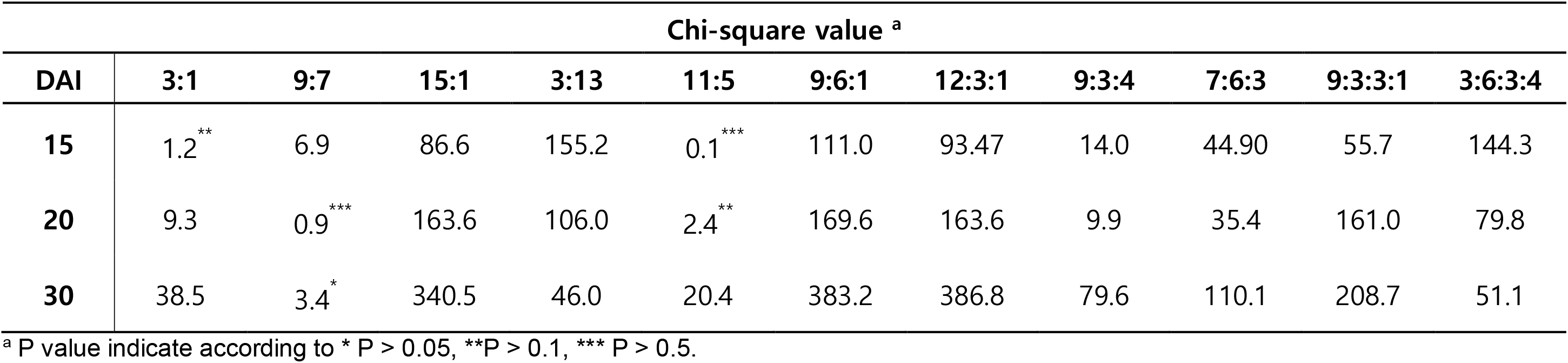
Segregation of *R. solanacearum* SL1931 resistance in F_2_ population at 15, 20, and 30 dai

| Chi-square value <sup>a</sup> |  |  |  |  |  |  |  |  |  |  |  |
| --- | --- | --- | --- | --- | --- | --- | --- | --- | --- | --- | --- |
| DAI | 3:1 | 9:7 | 15:1 | 3:13 | 11:5 | 9:6:1 | 12:3:1 | 9:3:4 | 7:6:3 | 9:3:3:1 | 3:6:3:4 |
| 15 | 1.2<sup>**</sup> | 6.9 | 86.6 | 155.2 | 0.1<sup>***</sup> | 111.0 | 93.47 | 14.0 | 44.90 | 55.7 | 144.3 |
| 20 | 9.3 | 0.9<sup>***</sup> | 163.6 | 106.0 | 2.4<sup>**</sup> | 169.6 | 163.6 | 9.9 | 35.4 | 161.0 | 79.8 |
| 30 | 38.5 | 3.4<sup>*</sup> | 340.5 | 46.0 | 20.4 | 383.2 | 386.8 | 79.6 | 110.1 | 208.7 | 51.1 |

<sup>a</sup> P value indicate according to \* P > 0.05, \*\*P > 0.1, \*\*\* P > 0.5.

## 3. Discussion

As global warming continues, the damage of BW is spreading beyond tropical and subtropical regions worldwide. The interaction between *R. solanacearum* and its plant hosts has been studied as plant resistance to bacterial phytopathogens for more than two decades [20, 37, 38]. To study various interactions with plants, it is important to establish accurate screening. Accordingly, the inoculation method that makes good use of the infection characteristic of the bacteria was dominated since *R. solanacearum* is a soil-dwelling bacterium. Soil-drench or/and root-dipping inoculation is mostly used to investigate bacterial wilt disease progress on peppers, tomatoes, eggplants, potatoes, and the model plants Medicago and Arabidopsis [15, 35, 37, 39–41]. Using this root-infection method requires a wound of the root, however, there is uncertainty regarding the infections before the symptoms alongside difficulty in knowing the exact resistance phenotype depending on the degree of artificial root wound. Consequentially, variation and deviation of the BW symptom appear large in plants [15–18]. To overcome these problems, we developed an LWB assay for BW on peppers.

In this study, we confirmed the different symptoms in leaves after inoculation to discover if the method is suitable for resistant ‘MC4’ and susceptible ‘Subicho’. Additionally, the transcript levels of defense-related genes and bacterial cell growth were significantly different in the resistant or susceptible cultivars following *R. solanacearum* infection. Although the strains and cultivars are different from our study, the result is consistent with the real-time visualization of the bioluminescent *R. solanacearum* strain BL-Rs7 colonization of grafted peppers in Du et al. (2019) that demonstrated more aggregation of the pathogen in susceptible cultivar (BVRC 1) then resistance (BVRC 25) [28]. Likewise, in our study, ‘MC4’ inhibited the proliferation of *R. solanacearum* and displayed a higher expression level of cell-death related genes compared with ‘Subicho’. The cell-death markers used in this study have related to the resistant response and defense-related pathway [32, 42]. As a result, it can be assumed that the resistance-related factor acts for the defense as ‘MC4’ has a higher expression value than that of ‘Subicho’. Through these results, we confirmed the ‘MC4’ was a clear BW resistance cultivar compared with 'Subicho'. According to the study of Akinori et al. (2007), the same BW phenotype was also represented in tobacco when leaf-infiltration and root-inoculation were performed, similar to our studies [43]. The leaf-infiltration method is more useful to elucidate molecular events than root (soil)-drenching to better understand the interaction between plants and pathogens since it is possible to inoculate equally [43–45]. In conclusion, the wilting symptoms appeared on the whole plant even when inoculated to the leaves, which was confirmed the same symptoms as the root infection.

The temperature is the main environmental factor in which *R. solanacearum* affects crops [46, 47]. An experiment was conducted to confirm the most suitable temperature conditions for LWB before the inoculum concentration experiment. As a result of our experiments at 25 °C, 28 °C, and 32°C, two suitable temperatures were revealed except for 25 °C (data not shown). Additionally, the studies derived that the temperature of 25 °C was not suitable for peppers and tomatoes, respectively, in the screening research for optimization condition [15, 35]. Therefore, the temperature was fixed at 28–30 °C in the experimental conditions, and the inoculum concentrations were tested to identify the most suitable for the LWB. The most appropriate concentration was 10^6^ CFU/mL indicating that it was sufficiently able to confirm the phenotypic difference between two control cultivars with a lower concentration and less volume than the drenching method.

We executed the LWB in eleven commercial pepper cultivars with BW phenotype information and one commercial pepper cultivar with no information. As a result, five and two cultivars represented R-phenotype and S-phenotype, respectively, and the others were the MR-phenotype. Among them, the cultivars of ‘Muhanjilju’, ‘PR-Jangwongeubje’, and ‘PR-Gukgadeapyo’ demonstrated susceptibility in Hwang et al. (2017), but our results demonstrated the resistance of BW phenotypes, which is an opposite result. These results could affect the metabolic activity of the host due to artificial wounds in the root, making it difficult to identify the accurate BW phenotype. In case of ‘Muhanjilju’ was represented R, MR, and S-phenotypes according to inoculation with various *R. solanacearum* strains in Lee et al. (2018) [16]. Additionaly, the ‘Subicho’ was inoculated by soil-drenching without root wounds and represented 0.7 DSI scores (0 to 4 scale scores) with very low disease incidence 15 dai [15], in which the BW phenotype is dependent on the root wound in pepper. For this reason, the study of interactions with pepper-*R. solanacearum* is exceptionally difficult. An accurate and reliable bioassay (LWB) can identify the exact BW phenotype in pepper through the equal inoculate without any wound of the root.

One of the most effective control managements is developing a resistance cultivar in the crops by integrating a resistance gene. Until now, a few sources of BW resistance have been reported in Capsicum spp. including *C. annuum* ‘MC4’, ‘MC5’, ‘LS2341’, and ‘PBC631’ [22-24]. In previous studies on the resource of resistance to BW, different QTL studies for only a few were determined that a major QTL (*qRRs-10.1*) in ‘BVRC1’ accession and one major (*Bw1*) in ‘LS2341’ accession were identified at different chromosome 10 and 1 for each resource, respectively [27, 28]. Despite the above reports of resistance to bacterial wilt, there are no useful cultivars comprised of high resistance with good yield and desirable agronomic traits. In this regard, understanding the genetic control for resistance to BW disease in plant breeding programs is essential and required to increase their efficiency, especially for planning a proper breeding method [48, 49].

‘MC4’ is well-known to have high-level resistance to the species of the *R. solanacearum* complex [22, 24, 29], but the genetic inheritance of ‘MC4’ for BW resistance has not been identified yet. In this study, we constructed the F_2_ population with ‘Subicho’ (susceptible) and performed an analysis of the inheritance of BW resistance through the LWB. We identified BW resistance was dominant over susceptible, and at least two pairs of genes appeared to control the trait in a complementary manner. Matsunaga et al. (1998) studied the mode of inheritance of BW resistance by crossing the resistant sweet pepper cultivar ‘Mie-Midori’ with the susceptible ‘AC2258’ and found that bacterial wilt resistance demonstrates incomplete dominance, and at least two genes were involved in resistance [25]. This result is similar to our segregation ratio date representing two major genes affected in the BW resistance of ‘MC4’ in this study. Additionally, Denis et al. (2005) concluded that two to five genes with additive effects were estimated to control the resistance. Tran et al. (2010) reported various dominance genetic effects as polygenic or oligogenic for *R. solanacearum* using 6 resistant pepper lines and 5 susceptible pepper lines [50]. Recently, Heshan et al. (2019) represented the disease index and wilt rate (%) using the F_2_ plants (n = 440), in which the wilting pattern of segregation was similar to our result [28]. Especially, the disease symptoms kept progressing over time alongside no represented complete dominance resistance like ‘MC4’ (R-parent). In the F_1_ and F_2_ generation, and which indicated to appear epistasis dominant like our result. These studies indicated that the inheritance of BW resistance is complicated, and a minimum of two genes interact to express resistance traits in the pepper germplasm. Our data suggest that the LWB method may determine a more exact BW resistance phenotype of pepper germplasms and reveal the interaction of plant-pathogens at the molecular level. Further investigations of inheritance factors could provide insights into QTL analysis and the development of BW resistance-related molecular markers.

## 4. Materials and Methods

### 4.1 Plant materials and growth conditions

Two varieties of peppers, *Capsicum annuum* ‘MC4’ with resistance to *R. solanacearum* and *C. annuum* ‘Subicho’ with susceptibility to *R. solanacearum*, were provided by Dr. Seon-Woo Lee (Dong-A University, Korea). The 12 commercial pepper cultivars (5 resistant, 5 moderately resistant, and 2 susceptible cultivars; Table 1) were used. The ‘MC4’ was crossed with ‘Subicho’ to get F_1_ plants. The F_2_ population was obtained by self-pollination of F_1_ plants. The pepper plants were kept in a growth chamber at 29 ± 1 °C under a 16 h light /8 h dark cycle with 50% humidity for 3–4 weeks. We inoculated *R. solanacearum* onto the 3rd and 4th leaves of fully expanded four-leaf-stage on pepper plants.

### 4.2 Bacteria inoculation and quantification

The strain *R. solanacearum* SL1931 (race1, phylotype I) was obtained from Dr. Seon-Woo Lee (Dong-A University, Korea). Bacterial cells were streaked and grown on Kelman’s tetrazolium chloride gar medium and maintained at 28 °C for 48-h. A single fluidal colony of *R. solanacearum* was grown on CPG broth and shaken at 250 rpm at 28 °C for 24-h. A bacterial culture suspension was diluted with distilled water to adjust the concentration to 10^8^ CFU/mL (OD_600_ = 0.3) [15]. Ten-fold serial dilutions of bacteria from 10^3^ CFU/mL to 10^6^ CFU/mL per leaf were used for inoculation. Seedlings at fully expanded four-leaf-stage were inoculated with 0.1 mL bacteria/leaf using a needleless syringe. Disease symptoms were observed under controlled conditions of 29 ± 1 °C under 16-h of light a day with 50% humidity for 20 days. The leaf-inoculation assay was performed with three independent tests, and each consisted of at least 8 plants per cultivar. Inoculum concentration was performed with 10^6^ CFU/mL per leaf for the inheritance analysis of the F_2_ population.

Bacterial quantification was performed like below with modification described by Yi et al. (2009) [51]. To determine in planta bacterial growth, pepper plants (*C. annuum* ‘MC4’ and ‘Subicho’) were leaf-inoculated with bacterial suspensions (1 × 10^4^ CFU/mL). Inoculated leaves were harvested at various time points for further analysis. Two independent assays were performed, which consisted of 6–8 samples for each time point in an experiment. Bacterial growth was measured by grinding inoculated samples in distilled water, plating serially diluted tissue samples with two replicates on CPG agar with 0.1% gentamicin (v/v), and counting colony-forming units.

### 4.3 Disease evaluation and data analysis

Disease evaluations were assessed daily after inoculation with *R. solanacearum* as described below. The disease severity index (DSI) of individual inoculated plants was rated on a scale of 0 to 4 as five phases in which 0 is no wilt disease symptoms observed; 1 is minor symptoms with less than 25% wilted leaves; 2 is moderate symptoms with 25–50% wilted leaves; 3 is severe symptoms with 50–75% wilted leaves; 4 is 75–100% wilted leaves or dead plant. The area under the disease progress curve (AUDPC) was calculated during the disease observation (0 to 15 dai) with a DSI value. [52]. Wilting rate (%) was calculated [The number of wilt plant / the number of total plants] x 100. The differences between the mean values of disease scores of the pepper cultivars were analyzed using Duncan’s multiple range tests, and p < 0.05 was considered a significant difference. Statistical analysis used SAS (SAS 9.1, SAS Institute Inc., Cary, NC, USA).

### 4.4 Quantitative RT-PCR of defense related genes

Total RNA was extracted from pepper leaves inoculated with the pathogen using the Trizol reagent (Invitrogen, Carlsbad, USA), and 2 ug of total RNA were reverse transcribed using Superscript IV (Invitrogen, Carlsbad, USA). To confirm the plant response against *R. solanacearum* infection, quantitative RT-PCR was performed using the defense-related genes (Supplementary Table S1) [32]. The following cycling conditions were used: 1 cycle of 94 °C for 3 min; 28 cycles or 30 cycles of 95 °C for 30 s, 58 °C for 30 s and 72 °C for 30 s; 72 °C for 5 min. The actin gene (designated *CaACT*) was used as an endogenous control to normalize the expression levels. Expression levels were reported as three replicates as mean values with standard errors.

## 5. Conclusions

Breeding a resistant cultivar is most effective in controlling bacterial wilt that causes serious yield losses in peppers worldwide. An accurate and reliable evaluation method is necessary to evaluate disease severity and reveal the genetic inheritance for BW resistance. We established a simple LWB to evaluate BW disease and then, using this, analyzed the inheritance of BW resistance through a ‘Subicho’ × ‘MC4’ F_2_ population. The BW resistance response of ‘MC4’ represents lower disease symptoms in leaves than susceptible ‘Subicho’, and we observed the spreading of wilt symptoms from leaves to a whole susceptible plant, similar to the drenching method. As a result, we optimized the evaluation method of resistance to BW with 12 commercial pepper cultivars. Using LWB, we confirmed the two major complementary genes related to the BW resistance trait through the analyzed genetic inheritance in 90 F_2_ progenies. This bioassay could promote an accurate evaluation of BW disease phenotype, and the two inheritance factors of ‘MC4’ could provide useful information for further QTL analysis in pepper breeding.

## Supplementary Materials

Supplementary Table S1. Primer information used for RT-PCR analysis of defense-related gene expression in this study. Supplementary Table S2. Disease evaluation design and the number of plants to parents and their progenies based on disease severity index in 15, 20, and 30 dai *against R. solanacearum* SL1931 strain.

## Funding

This research was supported by the Basic Science Research Program through the National Research Foundation of Korea (NRF) funded by the Korean Government (NRF-2017R1E1A1A01072843, 2015R1A6A1A03031413 and 2019R1C1C1007472). J.-S.K. were supported by a scholarship from the BK21 four Program from the Ministry of Education.

## Acknowledgments

We are grateful to Dr. Seon-Woo Lee (Dong-A University, Korea) for providing pepper seeds and bacterial strain.

## Author contributions

J.-S.K. performed the experiments, data analysis and wrote the manuscript. J.-Y.N. collected samples and data analysis. W.-H.K. and S.-I.Y. conceived and designed the experiments, organized and wrote the manuscript, and supervised the project.

## Institutional Review Board Statement

Not applicable.

## Informed Consent Statement

Not applicable.

## Conflicts of Interest

The authors declare no conflict of interest.

## Notes

### Competing Interest Statement

The authors have declared no competing interest.

## References

1. Howard, L. R.; Wildman, R. E., Antioxidant vitamin and phytochemical content of fresh and processed pepper fruit (*Capsicum annuum*). Handbook of nutraceuticals and functional foods. Boca Raton, 2007, pp. 165–191.

2. Faostat 2020. Available online: http://www.fao.org/ (accessed on 22 October 2020).

3. UN Comtrade Database. Abailable online: http://comtrade.un.org/ (accessed on 15 October 2020).

4. Common Names of Plant Disease. Available online : https://www.apsnet.org/edcenter/resources/commonnames/Pages/default.aspx (accessed on 24 October 2020).

5. Mansfield, J.; Genin, S.; Magori, S.; Citovsky, V.; Sriariyanum, M.; Ronald, P.; Dow, M.; Verdier, V.; Beer, S. V.; Machado, M. A., Top 10 plant pathogenic bacteria in molecular plant pathology. Molecular plant pathology 2012, 13, 614–629.

6. Jeong, Y.; Kim, J.; Kang, Y.; Lee, S.; Hwang, I., Genetic diversity and distribution of Korean isolates of *Ralstonia solanacearum*. Plant Disease 2007, 91, 1277–1287.

7. Lee, Y. K.; Kang, H. W., Physiological, biochemical and genetic characteristics of *Ralstonia solanacearum* strains isolated from pepper plants in Korea. Research in Plant Disease 2013, 19, 265–272.

8. Jiang, G.; Peyraud, R.; Remigi, P.; Guidot, A.; Ding, W.; Genin, S.; Peeters, N., Modeling and experimental determination of infection bottleneck and within-host dynamics of a soil-borne bacterial plant pathogen. bioRxiv 2016, 061408.

9. Jiang, G.; Wei, Z.; Xu, J.; Chen, H.; Zhang, Y.; She, X.; Macho, A. P.; Ding, W.; Liao, B., Bacterial wilt in China: history, current status, and future perspectives. Frontiers in Plant Science 2017, 8, 1549.

10. Hayward, A., Biology and epidemiology of bacterial wilt caused by *Pseudomonas solanacearum*. Annual review of phytopathology 1991, 29, 65–87.

11. Guidot, A.; Prior, P.; Schoenfeld, J.; Carrere, S.; Genin, S.; Boucher, C., Genomic structure and phylogeny of the plant pathogen *Ralstonia solanacearum* inferred from gene distribution analysis. Journal of bacteriology 2007, 189, 377–387.

12. Prior, P.; Ailloud, F.; Dalsing, B. L.; Remenant, B.; Sanchez, B.; Allen, C., Genomic and proteomic evidence supporting the division of the plant pathogen *Ralstonia solanacearum* into three species. BMC genomics 2016, 17, 90.

13. Safni, I.; Cleenwerck, I.; De Vos, P.; Fegan, M.; Sly, L.; Kappler, U., Polyphasic taxonomic revision of the *Ralstonia solanacearum* species complex: proposal to emend the descriptions of *Ralstonia solanacearum* and *Ralstonia syzygii* and reclassify current *R. syzygii* strains as *Ralstonia syzygii* subsp. syzygii subsp. nov., *R. solanacearum* phylotype IV strains as *Ralstonia syzygii* subsp. indonesiensis subsp. nov., banana blood disease bacterium strains as *Ralstonia syzygii* subsp. celebesensis subsp. nov. and *R. solanacearum* phylotype I and III strains as *Ralstonia pseudosolanacearum* sp. nov. International journal of systematic and evolutionary microbiology 2014, 64, 3087–3103.

14. Vasse, J.; Frey, P.; Trigalet, A., Microscopic studies of intercellular infection and protoxylem invasion of tomato roots by *Pseudomonas solanacearum*. Molecular Plant-Microbe Interactions 1995, 8, 241–251.

15. Hwang, S. M.; Jang, K. S.; Choi, Y. H.; Kim, H.; Choi, G. J., Development of an Efficient Bioassay Method to Evaluate Resistance of Chili Pepper Cultivars to *Ralstonia solanacearum*. Research in Plant Disease 2017, 23, 334–347.

16. Lee, J.; Lee, J.; Oh, D., Resistance of pepper cultivars to *Ralstonia solanacearum* isolates from major cultivated areas of chili peppers in Korea. Horticultural Science and Technology 2018, 36, 569–576.

17. Lee, H. J.; Jo, E. J.; Kim, N. H.; Chae, Y.; Lee, S. W., Disease responses of tomato pure lines against *Ralstonia solanacearum* strains from Korea and susceptibility at high temperature. Research in Plant Disease 2011, 17, 326–333.

18. Jung, E. J.; Joo, H. J.; Choi, S. Y.; Lee, S. Y.; Jung, Y. H.; Lee, M. H.; Kong, H. G.; Lee, S. W., Resistance evaluation of tomato germplasm against bacterial wilt by *Ralstonia solanacearum*. Research in Plant Disease 2014, 20, 253–258.

19. Fonseca, N. R.; Oliveira, L. S.; Guimarães, L. M.; Teixeira, R. U.; Lopes, C. A.; Alfenas, A. C., An efficient inoculation method of *Ralstonia solanacearum* to test wilt resistance in Eucalyptus spp. Tropical Plant Pathology 2016, 41, 42–47.

20. Huet, G., Breeding for resistances to *Ralstonia solanacearum*. Frontiers in plant science 2014, 5, 715.

21. Lee, S. M.; Kwak, Y. S.; Lee, K. H.; Kim, H. T., Control efficacy of fungicides on pepper bacterial wilt. The Korean Journal of Pesticide Science 2015, 19, 323–328.

22. Lopes, C. A.; Boiteux, L. S., Biovar-specific and broad-spectrum sources of resistance to bacterial wilt (*Ralstonia solanacearum*) in *Capsicum*. Embrapa Hortaliças-Artigo em periódico indexado (ALICE) 2004.

23. Mimura, Y.; Yoshikawa, M.; Hirai, M., Pepper accession LS2341 is highly resistant to *Ralstonia solanacearum* strains from Japan. HortScience 2009, 44, 2038–2040.

24. Tran, N. H.; Kim, B. S., Sources of resistance to bacterial wilt found in Vietnam collections of pepper (*Capsicum annuum*) and their nuclear fertility restorer genotypes for cytoplasmic male sterility. Plant Pathol. J 2012, 28, 418–422.

25. Matsunaga, H.; Sato, T.; Monma, S. In Inheritance of bacterial wilt resistance in the sweet pepper cv. Mie-Midori. In Proceeding of the 10th Eucarpia Meeting on Genetics and Breeding of Capsicum and Eggplant, Avignon, France, 7–11 Sep 1998; p. 172.

26. Lafortune, D.; Béramis, M.; Daubèze, A. M.; Boissot, N.; Palloix, A., Partial resistance of pepper to bacterial wilt is oligogenic and stable under tropical conditions. Plant Disease 2005, 89, 501–506.

27. Mimura, Y.; Kageyama, T.; Minamiyama, Y.; Hirai, M., QTL analysis for resistance to *Ralstonia solanacearum* in *Capsicum* accession ‘LS2341’. Journal of the Japanese Society for Horticultural Science 2009, 78, 307–313.

28. Du, H.; Wen, C.; Zhang, X.; Xu, X.; Yang, J.; Chen, B.; Geng, S., Identification of a major QTL (*qRRs-10.1*) that confers resistance to *Ralstonia solanacearum* in pepper *(Capsicum annuum*) using SLAF-BSA and QTL mapping. International journal of molecular sciences 2019, 20, 5887.

29. Lebeau, A.; Daunay, M. C.; Frary, A.; Palloix, A.; Wang, J. F.; Dintinger, J.; Chiroleu, F.; Wicker, E.; Prior, P., Bacterial wilt resistance in tomato, pepper, and eggplant: genetic resources respond to diverse strains in the *Ralstonia solanacearum* species complex. Phytopathology 2011, 101, 154–165.

30. Kim, B.; Cheung, J.; Cha, Y.; Hwang, H. Resistance to bacterial wilt of introduced peppers. 1998, 14, 217–219

31. Yi, S. Y.; Lee, D. J.; Yeom, S. I.; Yoon, J.; Kim, Y. H.; Kwon, S. Y.; Choi, D., A novel pepper (*Capsicum annuum*) receptor-like kinase functions as a negative regulator of plant cell death via accumulation of superoxide anions. New Phytol 2010, 185, 701–15.

32. Simko, I.; Piepho, H. P., The area under the disease progress stairs: calculation, advantage, and application. Phytopathology 2012, 102, 381–389.

33. Yeom, S. I.; Baek, H. K.; Oh, S. K.; Kang, W. H.; Lee, S. J.; Lee, J. M.; Seo, E.; Rose, J. K.; Kim, B. D.; Choi, D., Use of a secretion trap screen in pepper following *Phytophthora capsici* infection reveals novel functions of secreted plant proteins in modulating cell death. Molecular plant-microbe interactions 2011, 24, 671–684.

34. Huh, S. U.; Kim, K. J.; Paek, K. H., *Capsicum annuum* basic transcription factor 3 (*CaBtf3*) regulates transcription of pathogenesis-related genes during hypersensitive response upon *Tobacco mosaic virus* infection. Biochemical and biophysical research communications 2012, 417, 910–917.

35. Hamada, H.; Takeuchi, S.; Kiba, A.; Tsuda, S.; Suzuki, K.; Hikichi, Y.; Okuno, T., Timing and extent of hypersensitive response are critical to restrict local and systemic spread of *Pepper mild mottle virus* in pepper containing the *L3* gene. Journal of General Plant Pathology 2005, 71, 90–94.

36. Kim, B.; Hwang, I. S.; Lee, H. J.; Lee, J. M.; Seo, E.; Choi, D.; Oh, C. S., Identification of a molecular marker tightly linked to bacterial wilt resistance in tomato by genome-wide SNP analysis. Theoretical and Applied Genetics 2018, 131, 1017–1030.

37. Lee, J. H.; Jang, K. S.; Choi, Y. H.; Kim, J. C.; Choi, G. J., Development of an efficient screening system for resistance of tomato cultivars to *Ralstonia solanacearum*. Research in Plant Disease 2015, 21, 290–296.

38. Bocsanczy, A. M.; Achenbach, U. C.; Mangravita N, A.; Yuen, J. M.; Norman, D. J., Comparative effect of low temperature on virulence and twitching motility of *Ralstonia solanacearum* strains present in Florida. Phytopathology 2012, 102, 185–194.

39. Deslandes, L.; Pileur, F.; Liaubet, L.; Camut, S.; Can, C.; Williams, K.; Holub, E.; Beynon, J.; Arlat, M.; Marco, Y., Genetic characterization of *RRS1*, a recessive locus in *Arabidopsis thaliana* that confers resistance to the bacterial soilborne pathogen *Ralstonia solanacearum*. Molecular Plant-Microbe Interactions 1998, 11, 659–667.

40. Aoun, N.; Tauleigne, L.; Lonjon, F.; Deslandes, L.; Vailleau, F.; Roux, F.; Berthomé, R., Quantitative disease resistance under elevated temperature: genetic basis of new resistance mechanisms to *Ralstonia solanacearum*. Frontiers in plant science 2017, 8, 1387.

41. Lebeau, A.; Gouy, M.; Daunay, M. C.; Wicker, E.; Chiroleu, F.; Prior, P.; Frary, A.; Dintinger, J., Genetic mapping of a major dominant gene for resistance to *Ralstonia solanacearum* in eggplant. Theoretical and Applied Genetics 2013, 126, 143–158.

42. Cruz, A. P. Z.; Ferreira, V.; Pianzzola, M. J.; Siri, M. I.; Coll, N. S.; Valls, M., A novel, sensitive method to evaluate potato germplasm for bacterial wilt resistance using a luminescent *Ralstonia solanacearum* reporter strain. Molecular Plant-Microbe Interactions 2014, 27, 277–285.

43. Wang, K.; Remigi, P.; Anisimova, M.; Lonjon, F.; Kars, I.; Kajava, A.; Li, C. H.; Cheng, C. P.; Vailleau, F.; Genin, S., Functional assignment to positively selected sites in the core type III effector *RipG7* from *Ralstonia solanacearum*. Molecular plant pathology 2016, 17, 553–564.

44. Pontier, D.; Godiard, L.; Marco, Y.; Roby, D., *Hsr203J*, a tobacco gene whose activation is rapid, highly localized and specific for incompatible plant/pathogen interactions. Plant J 1994, 5, 507–21.

45. Kiba, A.; Maimbo, M.; Kanda, A.; Tomiyama, H.; Ohnishi, K.; Hikichi, Y., Isolation and expression analysis of candidate genes related to *Ralstonia solanacearum*–tobacco interaction. Plant Biotechnology 2007, 24, 409–416.

46. Nakano, M.; Nishihara, M.; Yoshioka, H.; Takahashi, H.; Sawasaki, T.; Ohnishi, K.; Hikichi, Y.; Kiba, A., Suppression of *DS1* phosphatidic acid phosphatase confirms resistance to *Ralstonia solanacearum* in *Nicotiana benthamiana*. PLoS One 2013, 8, e75124.

47. Planas M. M.; Bernardo F. M.; Paulus, J.; Kaschani, F.; Kaiser, M.; Valls, M.; van der Hoorn, R. A.; Coll, N. S., Protease activities triggered by *Ralstonia solanacearum* infection in susceptible and tolerant tomato lines. Molecular & Cellular Proteomics 2018, 17, 1112–1125.

48. Janse, J.; Van den Beld, H.; Elphinstone, J.; Simpkins, S.; Tjou-Tam-Sin, N.; Van Vaerenbergh, J., Introduction to Europe of *Ralstonia solanacearum* biovar 2, race 3 in *Pelargonium zonale* cuttings. Journal of Plant Pathology 2004, 147–155.

49. Singh, D.; Yadav, D.; Sinha, S.; Choudhary, G., Effect of temperature, cultivars, injury of root and inoculums load of *Ralstonia solanacearum* to cause bacterial wilt of tomato. Archives of Phytopathology and Plant Protection 2014, 47, 1574–1583.

50. Caranta, C.; Palloix, A., Both common and specific genetic factors are involved in polygenic resistance of pepper to several *potyviruses*. Theoretical and Applied Genetics 1996, 92, 15–20.

51. Thabuis, A.; Palloix, A.; Pflieger, S.; Daubeze, A. M.; Caranta, C.; Lefebvre, V., Comparative mapping of *Phytophthora* resistance loci in pepper germplasm: evidence for conserved resistance loci across Solanaceae and for a large genetic diversity. Theoretical and Applied Genetics 2003, 106, 1473–1485.

52. Tran, N. H.; Kim, B. S., Inheritance of resistance to bacterial wilt (*Ralstonia solanacearum*) in pepper (*Capsicum annuum* L.). HORTICULTURE ENVIRONMENT and BIOTECHNOLOGY 2010, 51, 431–439.

53. Kim, H. J.; Baek, K. H.; Lee, S. W.; Kim, J.; Lee, B. W.; Cho, H. S.; Kim, W. T.; Choi, D.; Hur, C. G., Pepper EST database: comprehensive in silico tool for analyzing the chili pepper (*Capsicum annuum*) transcriptome. BMC Plant Biology 2008, 8, 1–7.

54. Mateos, R. M.; Bonilla V., D.; Del Río, L. A.; Palma, J. M.; Corpas, F. J., *NADP*‐dehydrogenases from pepper fruits: effect of maturation. Physiologia Plantarum 2009, 135, 130–139.

