## Supplemental Table 1, 2 for "Leaf-to-whole plant spread bioassay for pepper and *Ralstonia solanacearum* interaction determines inheritance of resistance to bacterial wilt for further breeding"

**Supplementary Table S1.** Primer information used for RT-PCR analysis of cell death gene expression in this study.

| **Clone name** | **Sequence information (5'->3')** | | **Tm (^o^C)** | **size**  **(bp)** | **Accession No.**  **(NCBI)** | **References** |
| --- | --- | --- | --- | --- | --- | --- |
|  | **Forward** | **Reverse** |  |  |  |  |
| ***CaHsr203J*** | CGCCATCAGCGTCTCCGTCTTC | CGCCCATCGGACATGTGATTG | **58** | **413** | **AB162220** | **[****33****]** |
| ***CaCDM1*** | AAAGGCGCAAACGGGAGCTG | TCACAGCCAAACGCTTCGATCC | **60** | **400** | **GD056272** | **[****53****]** |
| ***CaHIN1*** | ATCTTCACTATTCTCATCGTCCTTG | TGCAAACTGCTAGTATTCTTGTGAC | **58** | **296** | **AB162221** | **[****33****]** |
| ***CaActin*** | CCACCTCTTCACTCTCTGCTCT | ACTAGGAAAAACAGCCCTTGGT | **58** | **165** | **AY572427** | **[****54****]** |

**Supplementary Table S2.** Disease evaluation design and the number of plants to parents and their progenies based on disease severity index in 15, 20, and 30 dai against *R. solanacearum* SL1931 strain.

| **Days after inoculation** | **Population ^a^** | **No. of Plants** | **Disease severity index** | | | | | **Mean of DSI ^b^** | **Wilt rate (%) ^c^** | **AUDPC ^d^** |
| --- | --- | --- | --- | --- | --- | --- | --- | --- | --- | --- |
|  |  |  | **0** | **1** | **2** | **3** | **4** |  |  |  |
| 15 | MC4 | 30 | 8 | 22 | 0 | 0 | 0 | 0.7 | 0 | 7.5 |
|  | Subicho | 30 | 0 | 0 | 0 | 0 | 30 | 4.0 | 100 | 50.3 |
|  | F_1_ | 30 | 0 | 17 | 2 | 1 | 10 | 2.13 | 36.6 | 22.7 |
|  | F_2_ | 90 | 0 | 60 | 3 | 7 | 20 | 1.89 | 32.2 | 21.9 |
| 20 | MC4 | 30 | 6 | 24 | 0 | 0 | 0 | 0.8 | 0 | 7.5 |
|  | Subicho | 30 | 0 | 0 | 0 | 0 | 30 | 4.0 | 100 | 50.3 |
|  | F_1_ | 30 | 0 | 12 | 4 | 0 | 14 | 2.5 | 46.7 | 22.7 |
|  | F_2_ | 90 | 0 | 44 | 11 | 1 | 34 | 2.3 | 38.8 | 21.9 |
| 30 | MC4 | 30 | 0 | 30 | 0 | 0 | 0 | 1.0 | 0 | 16.5 |
|  | Subicho | 30 | 0 | 0 | 0 | 0 | 30 | 4.0 | 100 | 90.3 |
|  | F_1_ | 30 | 0 | 0 | 7 | 1 | 22 | 3.5 | 73.3 | 52.9 |
|  | F_2_ | 90 | 0 | 1 | 41 | 3 | 46 | 3.0 | 54.4 | 48.5 |

^a^ ‘MC4’ and ‘Subicho’ is resistance (R) and susceptible (S) parent line, respectively. The F_1_ population crossed ‘Subicho’ (S) x ‘MC4’ (R) and F_2_ population derived from self-cross of F_1_ plants.
